# Quantitative in toto live imaging analysis of apical nuclear migration in the mouse telencephalic neuroepithelium

**DOI:** 10.1101/2024.09.25.614873

**Authors:** Tsukasa Shimamura, Takaki Miyata

**Affiliations:** Department of Anatomy and Cell Biology, Nagoya University Graduate School of Medicine, Nagoya, Japan. 65 Tsurumai, Showa, Aichi, Nagoya, Japan

**Keywords:** interkinetic nuclear migration, nuclear density, mean square displacement, tissue fluidity, Ripley’s K-function

## Abstract

In the embryonic neuroepithelium (NE), neural progenitor cells undergo cell cycle-dependent interkinetic nuclear migration (IKNM) along the apicobasal axis. Extensive IKNM supports increasing cell production rates per unit apical surface, as typically observed in the mammalian telencephalic NE. Apical nucleokinesis during the G2 phase is an essential premitotic event, but its occurrence has not yet been quantitatively analyzed at a large 3D scale with sufficient spatiotemporal resolution. Here, we comprehensively analyzed apically migrating nuclei/somata in reference to their surroundings from embryonic day (E)11 to E13 in the mouse telencephalon. The velocity of apical nucleokinesis decreased, with more frequent nuclear pausing occurring at E12 and E13, whereas the nuclear density in the middle NE zone (20-40 μm deep) increased. This result, together with the results of Shh-mediated overproliferation experiments in which the nuclear density was increased in vivo at E11, suggests that apical nucleokinesis is physically influenced by the surrounding nuclei. Mean square displacement analysis for nuclei being passed by the apically migrating nuclei via horizontal sectioning in toto-recorded movies revealed that the “tissue fluidity” or physical permissiveness of the NE to apical nucleokinesis gradually decreased (E11 > E12 > E13). To further investigate the spatial relationship between preexisting mitoses and subsequent premitotic apical nucleokinesis, the horizontal distribution of mitoses was cumulatively (∼3 hours) analyzed under in toto monitoring. The four-dimensional cumulative apical mitoses presented a “random”, not “clustered” or “regular”, distribution pattern throughout the period examined. These methodologies provide a basis for future comparative studies of interspecies differences.

## Introduction

In the embryonic neuroepithelium (NE), the neural progenitor cell (NPC) moves the nucleus to the basal side during the G1 phase of the cell cycle and then (after completion of DNA synthesis) returns to the apical surface during the G2 phase to undergo mitosis to produce daughter cells. This cell cycle-dependent nucleokinesis is called interkinetic nuclear migration (IKNM) (Sauer, 1935; Miyata et al., 2008; Norden, 2017), and its molecular mechanisms at the single-cell level have been explored in terms of actomyosin-dependent nuclear pushing/squeezing (Schenk et al., 2009, Spear and Erickson, 2012) and microtubule-dependent nuclear pulling (Tsai et al., 2005; Xie et al., 2007; Tsai et al., 2010; Cappello et al., 2011; Kosodo et al., 2011; Hu et al., 2013). Notably, NPCs perform their nucleokinetic and cell-production functions in a multicellular, three-dimensionally complex situation, in which different IKNM trajectories are laterally assembled in an asynchronous manner. Histologically, the appearance of heterogeneous nuclei in the cell cycle phase and migration status at different apicobasal positions of an epithelium are referred to as pseudostratification, the degree of which is high in NE, especially in the mammalian cerebral cortex (Sauer, 1935; Smart, 1972). Extensive pseudostratification is beneficial for increasing the cell-production rate per apical area in the epithelial system (Smart, 1972; Miyata et al., 2015). Failures in a certain phase of IKNM can secondarily result in cell community-level abnormalities such as overcrowding in the NE, leading to further pathological histogenesis (Okamoto et al., 2013). Evolutionally, the highly productive cerebral cortical NE in primates, including humans, is accompanied by extensive pseudostratification (Zecevic, 1993; Smart et al., 2002). Therefore, it is important to understand how the population-level IKNM is collectively organized to establish an ordered pattern.

As a basis for such understanding, comprehensive monitoring of the entire collective IKNM, i.e., how nuclei exclusively visualized across a full NE thickness behave in relation to their surroundings, is essential. We used mouse telencephalic NE, especially cerebral cortical NE, as a model system to investigate population-level IKNM. Given that basal nucleokinesis can be mechanically influenced (i.e., passively driven) by apical nucleokinesis (in zebrafish [Leung et al., 2011]; in mice [Kosodo et al., 2011; Shinoda et al., 2018]) and that apical nucleokinesis can be affected by the surroundings (in *Drosophila* [Kirkland et al., 2020; Hacht et al., 2022]), we sought to determine whether and, if so, how the apical nucleokinesis of the mouse cerebral NPC changes under possible influences by the surroundings, with a focus on the period from E11 to E13. This period is characterized by a consistently high mitotic frequency (Kowalczyk et al., 2009), whereas the density of NPCs peaks at E12 (Nishizawa et al., 2007), and the gene expression pattern in NPCs changes from proliferative to neurogenic toward E13 (Okamoto et al., 2016). Therefore, this period is attractive in that NPCs are consistently very active with extensive IKNM, whereas certain changes in the collective IKNM can be expected in relation to the transitions in cell production status.

## Materials and Methods

1. **Animals** R26-H2B-mCherry transgenic mice (accession No. CDB0239K; Abe et al., 2011) were provided by Toshihiko Fujimori (NIBB, Japan). Fucci mice (Fucci S/G2/M-Green#504, accession number RBRC02706; Sakaue-Sawano et al., 2008) were provided by Atsushi Miyawaki (RIKEN, Japan). ICR mice were purchased from Japan SLC. All the mice were housed under specific pathogen-free conditions at Nagoya University. The animal experiments were conducted according to the Japanese Act on Welfare and Management of Animals, Guidelines for Proper Conduct of Animal Experiments (published by the Science Council of Japan), and the Fundamental Guidelines for Proper Conduct of Animal Experiments and Related Activities in Academic Research Institutions (published by the Ministry of Education, Culture, Sports, Science, and Technology, Japan). All protocols for the animal experiments were approved by the Institutional Animal Care and Use Committee of Nagoya University (No. 29006). E0 was defined as the day of vaginal plug identification.
2. **Live observation of NPCs and the cerebral wall** Mouse embryos (E11-E13) isolated from the uterus were transferred to a Petri dish containing phosphate-buffered saline (PBS), and the head region that was surgically isolated was transferred to a dish containing DMEM/Ham’s F-12 culture medium. For coronal sectional observations, which enabled IKNM observation along the apicobasal axis, the brains were sliced (0.4 mm thick) via a vibratome. For observations in planes parallel to the apical surface (i.e., en face and horizontal sectional observations), the dorsal telencephalic wall was surgically trimmed to obtain a rectangle (0.5-1 mm on a side). These slices or rectangles of telencephalic walls were embedded in a polystyrene cell culture dish (Corning) with CellMatrix type-I collagen gel (Nitta Gelatin) at a concentration of 0.9–1.1 mg/ml and cultured in DMEM/Ham’s F-12 medium supplemented with 5% horse serum, 5% fetal bovine serum, N2 (1:100, Thermo Fisher Scientific) and B27 (1:50, Thermo Fisher Scientific) with 95% O_2_ and 5% CO_2_ at 37 °C during image acquisition. For imaging of all cells in the NE, the telencephalic walls were preincubated in DMEM/Ham’s F-12 medium supplemented with FM4-64 (Thermo Fisher Scientific) at a concentration of 5 μg/ml for 30 min, as described previously (Okamoto et al., 2014). Confocal time-lapse images were obtained with two systems as follows: a BX51W1 microscope (time intervals: 5 min, Z intervals: 2 μm) (Olympus) equipped with a CSU-X1 laser scanning confocal imaging unit (Yokogawa), a 40× objective lens (LUMPLFLN40XW, Olympus) or 60× objective lens (LUMFLN60XW, Olympus), an iXon+ EMCCD camera (Andor), an on-stage culture chamber (Tokai Hit), and a CV1000 laser scanning confocal imaging system (time intervals: 10 min, Z intervals: 1 mm) (Yokogawa) with a 40× objective lens (UPLFLN40XD, Olympus). Two-photon excitation images were obtained with an A1RMP (time intervals: 6 min, Z intervals: 2 μm) (Nikon) equipped with a Mai Tai DeepSee (Spectra Physics), a 25× objective lens (CFI75 APO 25XCW, Nikon) and an on-stage culture chamber (Tokai Hit), as described previously (Kawasoe et al., 2020). The excitation wavelength of FM4-64 was 950 nm.
3. **In utero electroporation and Immunohistochemistry** To achieve the overgrowth of NPCs in the cerebral wall, either pCAG-EGFP (0.5 mg/ml) or pCAGGS-Shh-N-ires-EGFP (1.0 mg/ml, gift from Jun Motoyama) (Shikata et al., 2011) was transfected into the telencephalon at E10 via in utero electroporation as described previously (Okamoto et al., 2013). The brains extracted at E11 were fixed in 4% paraformaldehyde/PBS for 1 h, immersed in 20% sucrose overnight, embedded in OCT compound (Sakura Finetek Japan), and then frozen. The frozen sections (16 mm thick) were incubated with an anti-BrdU antibody (BU1/75[ICR1], Abcam, 1/500) overnight at 4 °C. The sections were then treated with an Alexa546-conjugated secondary antibody (A-11081, Thermo Fisher Scientific, 1/500) with DAPI (Thermo Fisher Scientific, 1/1000) and subjected to confocal microscopy (FV1000, Olympus) with a 10x objective lens (UPlanSApo10X, Olympus) or 40x objective lens (UPlanFl40x, Olympus).
4. **Preprocessing of imaging data** The image sequences obtained via live observations were preprocessed via ImageJ/Fiji with plugins as follows: ‘Fast4Dreg’ (Laine et al., 2019) or ‘Correct 3D drift’ (Parslow et al., 2014) for the recursive alignment of the three-dimensional image stacks between time points, and ‘TransformJ’ for rotation and reslicing of the three-dimensional image stacks to obtain perpendicular or parallel planes to the apical surface (Meijering et al., 2001).
5. **Statistical analyses** Statistical analyses were performed via R for the Mann‒Whitney *U* test for two-group comparisons or the Steel‒Dwass test for three-group comparisons. All the statistical tests were two-tailed, and *p* < 0.05 was considered statistically significant. The actual *p* values are shown in each graph.
6. **Mean square displacement** The mean square displacement (MSD) is defined as:

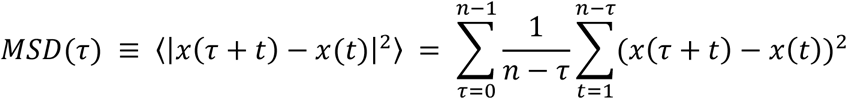

where:
  - τ represents the time lag.
  - x(t) is the position at time t.
  - n is the total number of time steps in the trajectory.
  This equation calculates the average squared displacement of a cell body over all possible time intervals of length τ. The angular brackets 〈…〉 denote an ensemble average. The first sum iterates over all possible time lags, from 0 to n-1. The second sum calculates the squared displacement for each pair of points separated by the time lag τ for all available pairs in the trajectory. The Factor 1/(n-τ) normalizes the inner sum, accounting for the decreasing number of available pairs as τ increases. For normal diffusion processes, the MSD has a linear relationship with time:

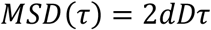

where:
  - d is the number of dimensions (e.g., d = 3 for three-dimensional space)
  - D is the diffusion coefficient.
  To obtain the diffusion coefficient D from the MSD data, we fit a straight line to the MSD plot and calculate the slope, m. The diffusion coefficient is then given by D = m/2d. To calculate lateral “tissue fluidity”, we set d to 2, and the diffusion coefficient was calculated from the 1-hour data.
7. **Spatiotemporal pattern analysis** We perform spatial analysis via Ripley’s K function, which calculates the statistical indicator K(r) from the spatial distribution pattern. This function was produced by the R package “spatstat” (Baddeley et al., 2015). The statistical indicator K(r) is defined as:

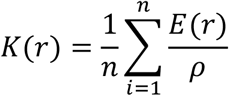

where:

- r represents the searching radius from an event.
- n is the number of total event points.
- where ρ is the event density in the total field.
- E(r) represents the number of events inside the circle radius = r.

In a random distribution pattern such as Poisson’s distribution, K(r) equals *πr*^2^. When analyzing the data, we perform 99 random pattern simulations and create a 95% random envelope, which contains 95% of the simulated lines. The observed event distribution data were then compared with the simulated 95% random envelope. If the observed line was above the random envelope, the observed distribution pattern was considered clustered, and if it was below, it was considered a regular (repelling-like) pattern.

## Results

### The apical nuclear migration speed decreased as the pausing time increased from E11 to E13

To comprehensively monitor the nuclei of apically migrating G2-phase NPCs, we first used Geminin-mAG reporter mice (consisting of the Fucci system) (Sakaue-Sawano et al., 2008), which exclusively visualize cells from the S phase to the early M phase (Figure 1a, 1b). We obtained 91 trajectories from E11 slices (n = 7), 71 trajectories from E12 slices (n = 7), and 70 trajectories from E13 slices (n = 6), in which the coordinates of the center of each apical migrating nucleus were recorded every 10 min, and the distance between the nucleus center and the apical surface, as well as the migration speed, was calculated. Although we could not directly distinguish S-phase NPCs from G2-phase NPCs, the beginning of apical migration, which is indicative of the entrance into the G2 phase (Kosodo et al.,2011), was readily recognized by detecting the generation and acceleration of the nuclear speed up to 10% of the maximum apical velocity. The total distance of apical nucleokinesis was not significantly different among E11 (42.3 ± 9.6 μm), E12 (44.9 ± 10.5 μm), and E13 (46.5 ± 11.3 μm) (p > 0.05 for each comparison) (Figure 1d). However, the time used for apical nucleokinesis was significantly shorter at E11 (95.7 ± 41.9 min) than at E12 (154.6 ± 62.5 min) and E13 (180.9 ± 90.9 min) (mean ± sd) (p < 0.001 between E11 and E12 and between E11 and E13) (Figure 1e). Consistently, the average speed during the entire period of apical nucleokinesis was significantly greater at E11 (29.9 ± 12.8 μm/min) than at E12 (20.2 ± 10.9 μm/min) and E13 (18.2 ± 8.6 μm/min) (mean ± sd) (p < 0.001 between E11 and E12 and between E11 and E13) (Figure 1f). Additionally, the maximum speed (10 min) was greater at E11 (61.3 ± 22.4 μm/min) than at E12 (52.4 ± 22.6 μm/min) and E13 (46.7 ± 19.3 μm/min) (mean ± sd) (p < 0.01 between E11 and E12 and p < 0.001 between E11 and E13), although the dominance of E11 over E12 or E13 decreased (Figure 1g). These results indicate that apical migration decelerates between E11 and later time points.

**Figure 1.**
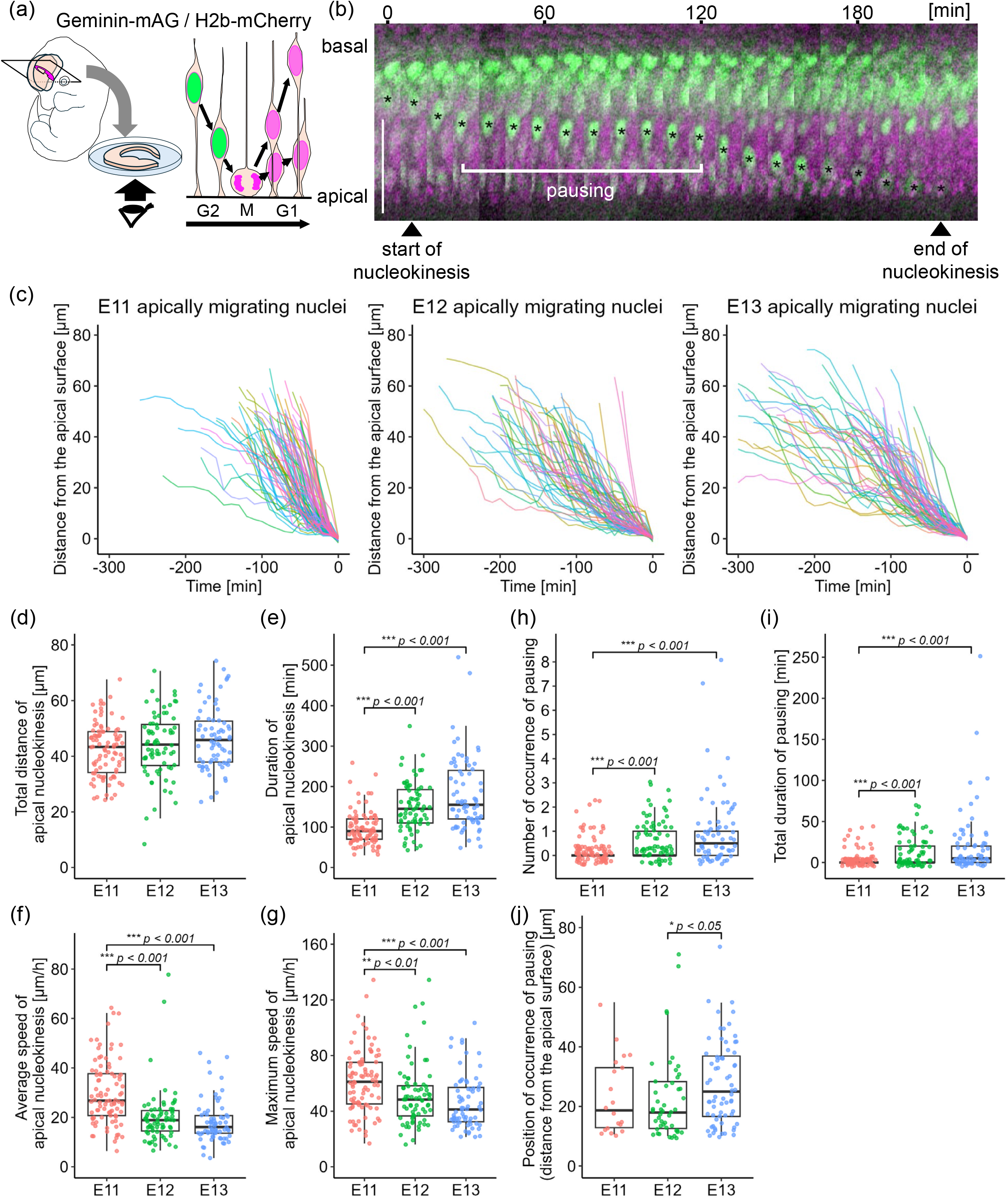
Comparison of live observed apical nucleokinesis during the G2 phase between embryonic days (E) 11, E12, and E13. (a) Schematic illustration depicting methods for coronal slicing of the embryonic telencephalon and imaging with the Geminin-mAG/H2b-mCherry system, in which the nuclei of S-G2-phase neural progenitor cells (NPCs) can be visualized in green. (b) Apical nucleokinesis shown by NPCs in an E12 slice. Scale, 50 μm. (c) Comparison of the trajectories of apically migrating nuclei. (d) Graph comparing the total distance of apical nucleokinesis. (e) Graph comparing the duration of apical nucleokinesis. (f) Graph comparing the average speed of apical nucleokinesis. (g) Graph comparing the maximum speed of apical nucleokinesis. (h) Graph comparing the number of nuclear pausings. (i) Graph comparing the total duration of nuclear pausing. (j) Graph comparing the position at which nuclear pausing occurred (the distance from the apical surface). Each comparison was performed via Steel–Dwass tests.

We observed that the apically migrating nuclei transiently slowed (to < 10% of the maximum speed) or stopped before they reached the apical surface (Figure 1b), which was referred to as pausing. Our comparison revealed that pausing during apical nucleokinesis was significantly more common at E12 (0.7 ± 0.1 times per apical nucleokinesis) and E13 (1.0 ± 0.2 times) than at E11 (0.2 ± 0.05 times) (mean ± se) (p < 0.001 between E11 and E12 and between E11 and E13) (Figure 1h). The total duration of pausing was also significantly longer at E12 (13.3 ± 2.2 min) and E13 (20.9 ± 4.8 min) than at E11 (4.1 ± 1.0 min) (mean ± se) (p < 0.001 between E11 and E12 and between E11 and E13) (Figure 1i). The position (i.e., the apicobasal depth) at which pausing occurred was almost comparable between E11 (23.3 ± 2.8 μm) and E12 (22.7 ± 2.0 μm), although pausing occurred at deeper levels at E13 (27.8 ± 1.7 μm) than at E12 (mean ± se) (p = 0.037 between E12 and E13) (Figure 1j).

### The density of telencephalic NE nuclei increased in the middle of the NE zone in a stage-dependent manner

To investigate the tissue-level mechanism potentially underlying the abovementioned deceleration of apical nucleokinesis with increased nuclear pausing after E11, we focused on the density of nuclei in the NE, especially the nuclear density in the middle zone of the NE (∼40 μm from the apical surface), across which G2-phase NPCs send nuclei to the apical surface. To this end, NE tissues prepared from H2B-mCherry mice (Abe et al., 2011), in which all cell nuclei (or condensed chromosomes in M-phase cells) were visualized, were horizontally sectioned (every 2-μm slice parallel to the apical surface) (Figure 2a, 2b). The number of nuclei per horizontal sectional area (0.1 mm × 0.1 mm) at 20 μm from the apical surface was significantly greater at E13 (204.7 ± 33.6) than at E11 (155.3 ± 18.2) and E12 (152.7 ± 13.5) (mean ± sd) (each p < 0.05). Similarly, the nuclear density at 30 μm was greater at E13 (219.2 ± 25.0) than at E11 (179.0 ± 12.8) and at E12 (177.7 ± 15.5) (mean ± sd) (each p < 0.05) (Figure 2c). We also measured the occupancy of a horizontal sectional area by nuclei at a depth of 30 μm and found that it gradually increased in a stage-dependent manner: 49.2% at E11, 54.2% at E12 (significantly greater than at E11 [p < 0.001], and 58.3% at E13 (significantly greater than at E12 [p < 0.001]) (Figure 2d). Notably, we observed pausing of apical nucleokinesis (Figure 1b) more frequently at E13 than at E11 or E12 in this middle NE zone (Figures 1h, 1i). This result is therefore consistent with the idea that the stage-dependent deceleration of apical nucleokinesis between E11 and later (Figures 1f, 1g) could be associated with the densification of nuclei in the NE, especially in a zone in which apically migrating G2-phase nuclei would need to physically share with stationary nuclei or basally migrating nuclei of non-G2-phase NPCs.

**Figure 2.**
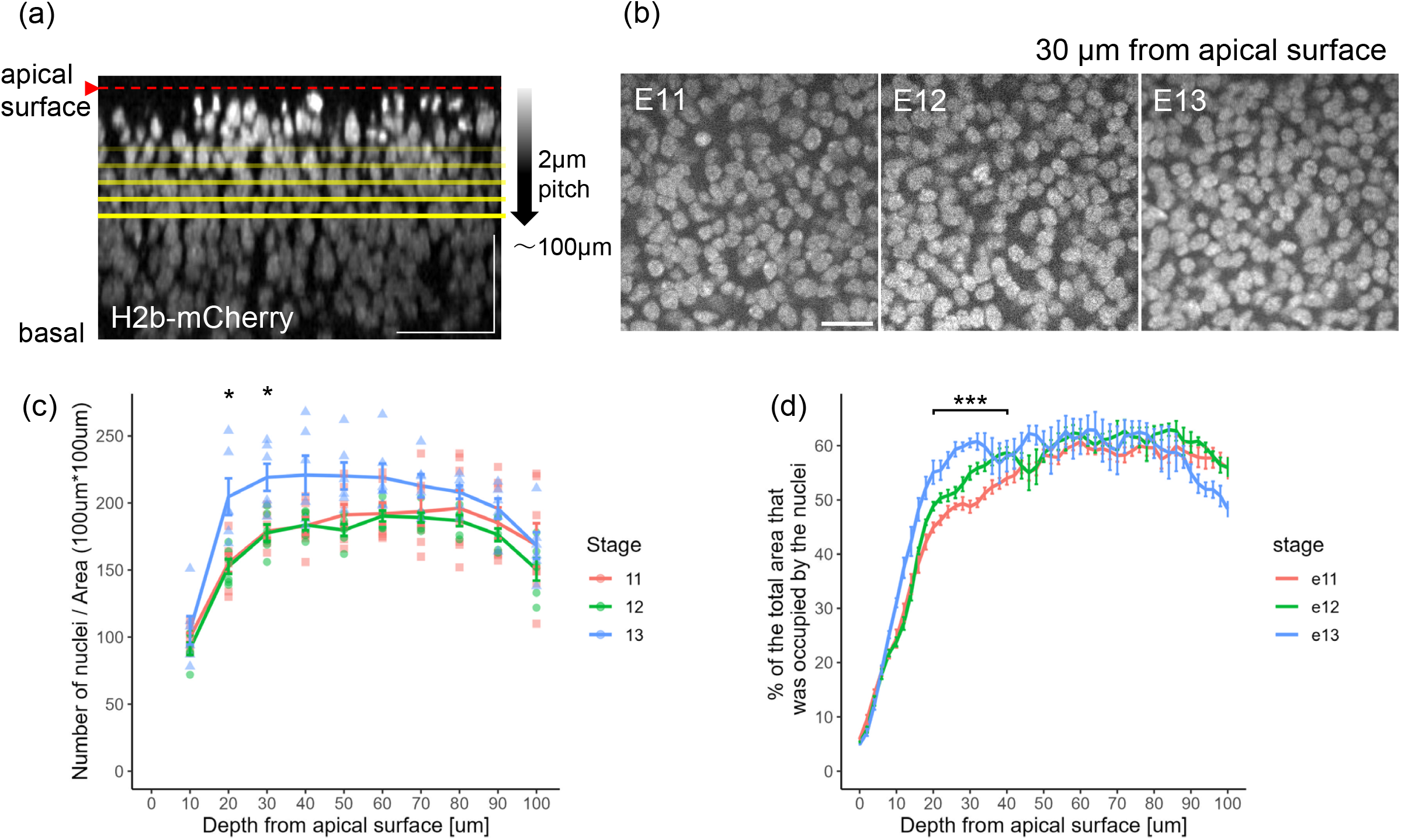
Comparison of the intra-NE nuclear density between E11, E12, and E13. (a) Method of measurement. Telencephalic walls prepared from H2B-mCherry mice were mounted on the apical top and observed via confocal microscopy at 2-μm intervals to 100-μm depth. (b) Horizontal section images taken at 30 μm from the apical surface. (c) Graph showing the depth-dependent changes in the density of nuclei (per 100 mm × 100 mm). (d) Graph showing the depth-dependent changes in the percentage of the total horizontal sectional area that was occupied by the nuclei. Scale bars: 50 μm (a) and 20 μm (b).

### Shh-mediated NE nuclear densification affected apical nucleokinesis in vivo

To experimentally test a model in which nuclear densification in the NE physically affects apical nucleokinesis, thereby leading to altered nuclear behaviors such as deceleration or pausing, we performed in utero electroporation (IUE) to introduce a Shh expression vector, which is known to induce the overproliferation of NPCs (Shikata et al., 2011). On the basis that nuclear density was lower at E11 than at E12 or E13, we aimed to determine, following IUE at E10, whether apical nucleokinesis in the Shh-overexpressing (Shh-OE) (and thus densified) E11 NE decelerated in a manner similar to that observed in the E12 or E13 NE tissues (Figure 3b). Low-power microscopic observations of frozen sections of the Shh-OE E11 cerebral walls revealed circumferential expansion/lengthening with frequent bucking (Figure 3c), which was considered to have occurred under confinement by the brain-surrounding contractile connective tissues (Tsujikawa et al., 2022) (Figure 3a). High-power examinations of the DAPI-stained sections revealed that the NE structure became thicker apicobasally (Figure 3c) in the Shh-treated group (126.7 ± 13.1 μm) than in the control group (92.6 ± 8.8 μm) (mean ± sd) (p < 0.01) and that the aspect ratio of the nucleus was greater in the Shh-OE group (2.5 ± 0.5) than in the control group (2.1 ± 0.4) (mean ± sd) (p < 0.001) (Figure 3d). These findings suggest that the Shh IUE induced lateral compression of the NE tissue. Similarly, the nuclear density within 40 mm from the apical surface was greater in the Shh-induced NE (116.9 ± 15.5) than in the control NE (107.2 ± 10.8) (p = 0.025) (Figure 3f). These findings suggest that Shh introduction effectively provided the physical conditions that we expected (Figure 3a). Unfortunately, however, direct assessment of apical nucleokinesis in this artificially densified E11 NE was not possible in culture. Surgical removal of such laterally compressed cerebral walls from the externally confined 3D environment resulted in a recoil of the wall in the circumferential direction. This release of the wall from the laterally compressed in vivo situation, thereby laterally loosening the NE to a normal nuclear density, precluded live monitoring under the required (NE densified) experimental conditions in culture.

**Figure 3.**
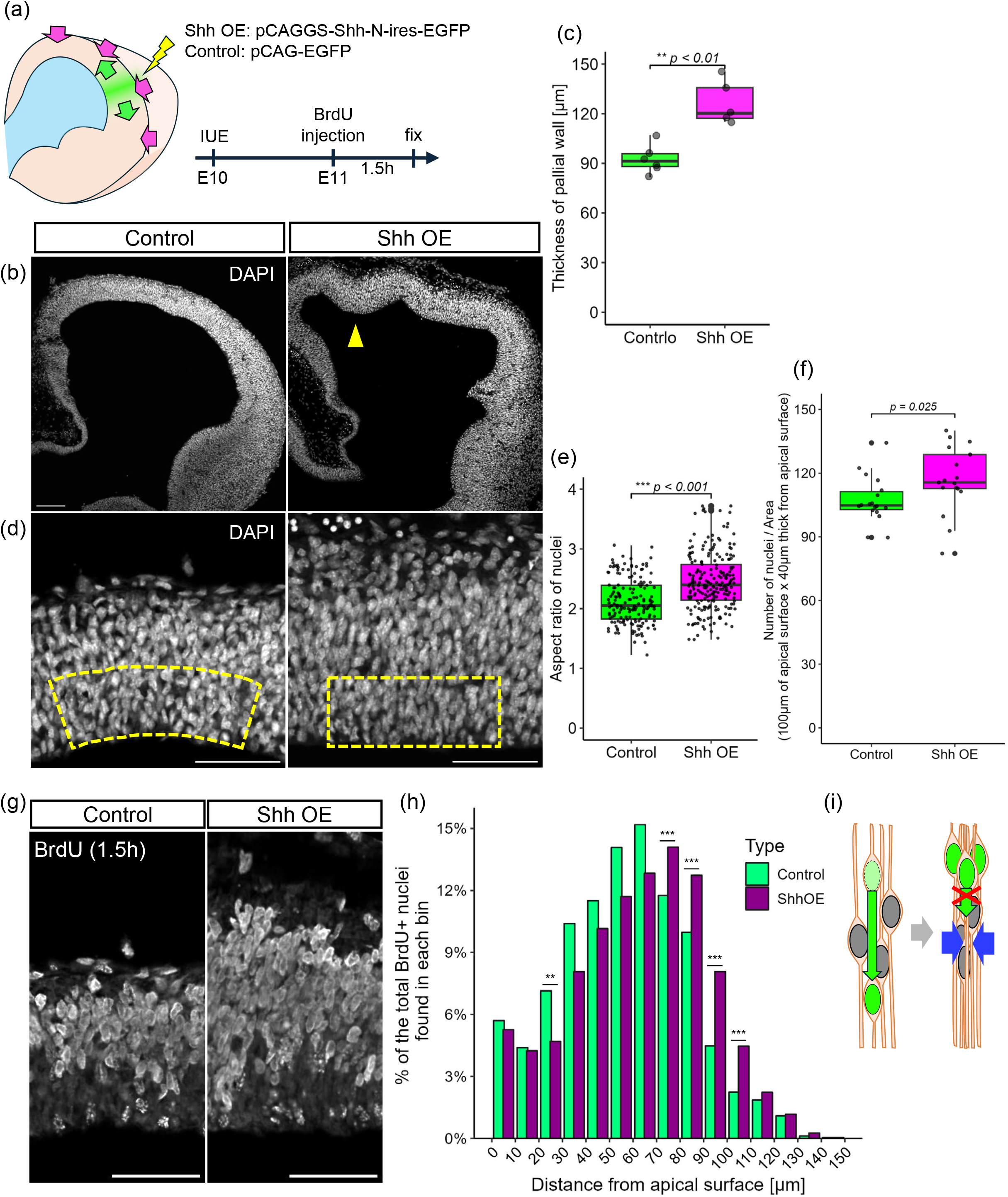
In vivo experiments were performed to assess the effect of Shh overexpression on intra-NE nuclear densification. (a) Telencephalic walls are expected to expand laterally under physical confinement from connective tissues, leading to tissue compression and nuclear densification. In utero electroporation (IUE) with a Shh vector at E10 was followed by bromodeoxyuridine (BrdU) injection at E11 1.5 hours before the embryos were fixed. (b) Photomicrographs comparing the control and Shh-overexpressing (OE) E11 telencephalic walls. Gray, DAPI. The arrowhead in the top (low-power) panel shows buckling of the wall. (c) Graph comparing the wall thickness between the control and Shh-OE groups. (d) High-magnification images of the control and Shh-OE telencephalic walls at E11. Gray, DAPI. (e) Graph comparing the aspect ratio of the nucleus (major axis/minor axis) between the control and Shh-OE groups. (f) Graph comparing the number of nuclei/area (nuclei number/40 μm thickness from the apical surface × 100 μm length of the apical surface) between the control and Shh-OE groups. (g) High-magnification images of the control and shh-OE telencephalic walls at E11, BrdU. (h) Histogram comparing the NE depth-dependent distribution of BrdU^+^ nuclei between the control and Shh-OE groups. (i) Schematic illustration of the physical restriction to apical nucleokinesis suggested in the Shh-induced nuclear densified NE. Scale bar: 100 μm (b), 50 μm (d, g).

Accordingly, we used an indirect (pulse-chase) method to assess apical nucleokinesis in vivo. Pulse administration of bromodeoxyuridine (BrdU) to E10-electroporated maternal mice at E11 was followed by fixation of the embryos 1.5 hours later. This in vivo protocol allows us to capture both a group of apically migrating nuclei of G2-phase NPCs (i.e., NPCs that have exited soon after BrdU administration) and another group of nuclei that are still in S phase (Takahashi et al., 1995; Hayes and Nowkowski, 2000) (Figure 3b). We compared the distribution of BrdU^+^ nuclei between the control and Shh-OE groups via a histogram to calculate the percentage of total BrdU^+^ nuclei observed in each bin (10 μm thick) (Figure 3g). The average distance between BrdU^+^ nuclei and the apical surface was significantly greater in the Shh-OE group (62.4 ± 28.4 μm) than in the control group (56.7 ± 27.1 μm) (mean ± sd) (p < 0.001) (Figure 3g). The histogram revealed that the four bins in the basal half of the NE (70-80 μm, 80-90 μm, 90-100 μm, and 100-110 μm) presented more BrdU^+^ nuclei in the Shh-OE compared with the control (p < 0.001 for each of the four basal bins, *χ*^2^ test), whereas the five bins (20-70 μm) apical to these basal bins presented fewer BrdU^+^ nuclei in the Shh-OE compared with the control (p < 0.01 in the 20-30 μm bin). Given that the DAPI-stained sections showed thickening of the NE in the Shh-OE group (126.7 μm) compared with those from the control group (92.6 μm) (Figure 3c), the former difference, i.e., an increase in the number of BrdU^+^ nuclei in the basal NE of the Shh-OE group, can be partly explained, especially in the bins 90-100 μm and 110-110 μm, in which the S-phase nuclei occupying the outer half of the NE (Takahashi et al., 1995; Hayes and Nowkowski, 2000) slightly shifted their overall distribution in the basal direction. The latter difference, i.e., a decrease in BrdU^+^ nuclei in the middle NE subzones of the Shh-OE group, especially in the 20-30 μm and 30-40 μm bins, can be almost solely regarded as G2-phase nuclei (Hayes and Nowkowski, 2000). Since apical nucleokinesis during the G2 phase can normally start from multiple intra-NE levels ranging from 30-70 μm deep (Hayes and Nowkowski, 2000), the possible Shh-induced basal shift of the basally located S-phase nuclei (i.e., from the 60-80 μm bin to the 80-100 μm bin) would not influence the distribution of BrdU^+^ nuclei in the 20-40 μm bins of the Shh-OE NE, allowing the reduced BrdU^+^ nuclei in the 20-40 μm bins to most strongly support the possibility that the apical nucleokinesis during the G2 phase was restricted to the Shh-OE NE. A group of G2-phase nuclei that should normally start apical nucleokinesis at 60-80 μm may also have been partly stacked at 80-100 μm. These results suggest that Shh-mediated NE densification affects apical nucleokinesis in vivo, supporting our model that apical nucleokinesis is affected physically by the surrounding environment.

### MSD analysis via horizontal section nuclear tracking suggested a stage-dependent change in the physical permissiveness of a 30-mm-deep zone to apical nucleokinesis

Why does increased nuclear density affect apical nucleokinesis? For the apically migrating nucleus of an NPC, the nuclei of other NPCs must be the major obstacle. Nevertheless, quantitative assessment of the ease or difficulty of passing through a certain space filled with cells has not yet been performed for NE. As an approach to realistically understand changes in the physical tissue status of NEs while increasing nuclear density in a stage-dependent manner, we used two-photon microscopically obtained horizontal sectional movies of NE tissues labeled with FM4-64, in which the cell membrane was comprehensively visualized (Kawaue et al., 2014). When the nucleus/soma of a non-G2-phase NPC is passed by an apically migrating G2-phase nucleus, the former (apicobasally stational) nucleus can be orthogonally (laterally) displaced (Figure 4a). In such a case, there was space for the former (apicobasally stational) nucleus to be pushed and drifted into, suggesting that this horizontal plane of the NE possesses a considerable level of physical permissiveness to the latter (apically migrating) nucleus. Conversely, if such space for lateral nuclear dodging is reduced in a crowded situation, apical nucleokinesis would physically encounter difficulty. Thus, objective assessment of horizontal nuclear drifts should be useful for connecting alterations in apical nucleokinesis with NE nuclear densification.

**Figure 4.**
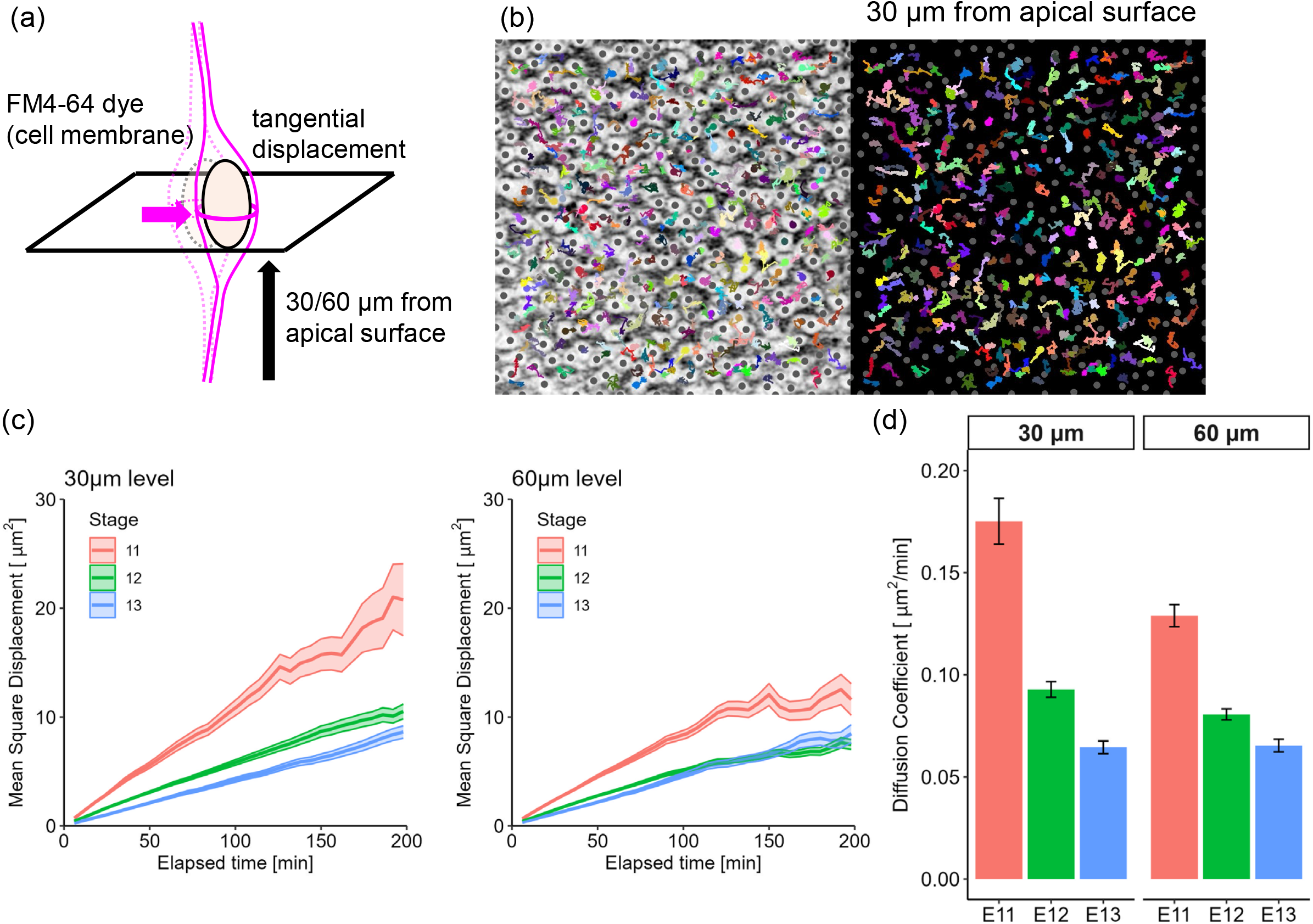
MSD analysis via horizontal section nuclear tracking to assess the physical permissiveness of an intra-NE subzone to apical nucleokinesis. (a) Tangential displacement of the nucleus/soma of each NPC in response to nearby apical nucleokinesis, schematically illustrated for analysis at the level of 30 or 60 μm from the apical surface. (b) Tracking (6 hours) of the horizontal nuclear drifts of individual nuclei/somata in NE (30 μm deep) superimposed with a photomicrograph (left) of horizontal sectionally obtained FM4-64-labeled cell‒cell borders (black). (c) Graph comparing the mean square displacement (MSD) plot between E11, E12, and E13. (d) Graph comparing the diffusion coefficients between E11, E12, and E13 and between 30 and 60 μm.

**Figure 5.**
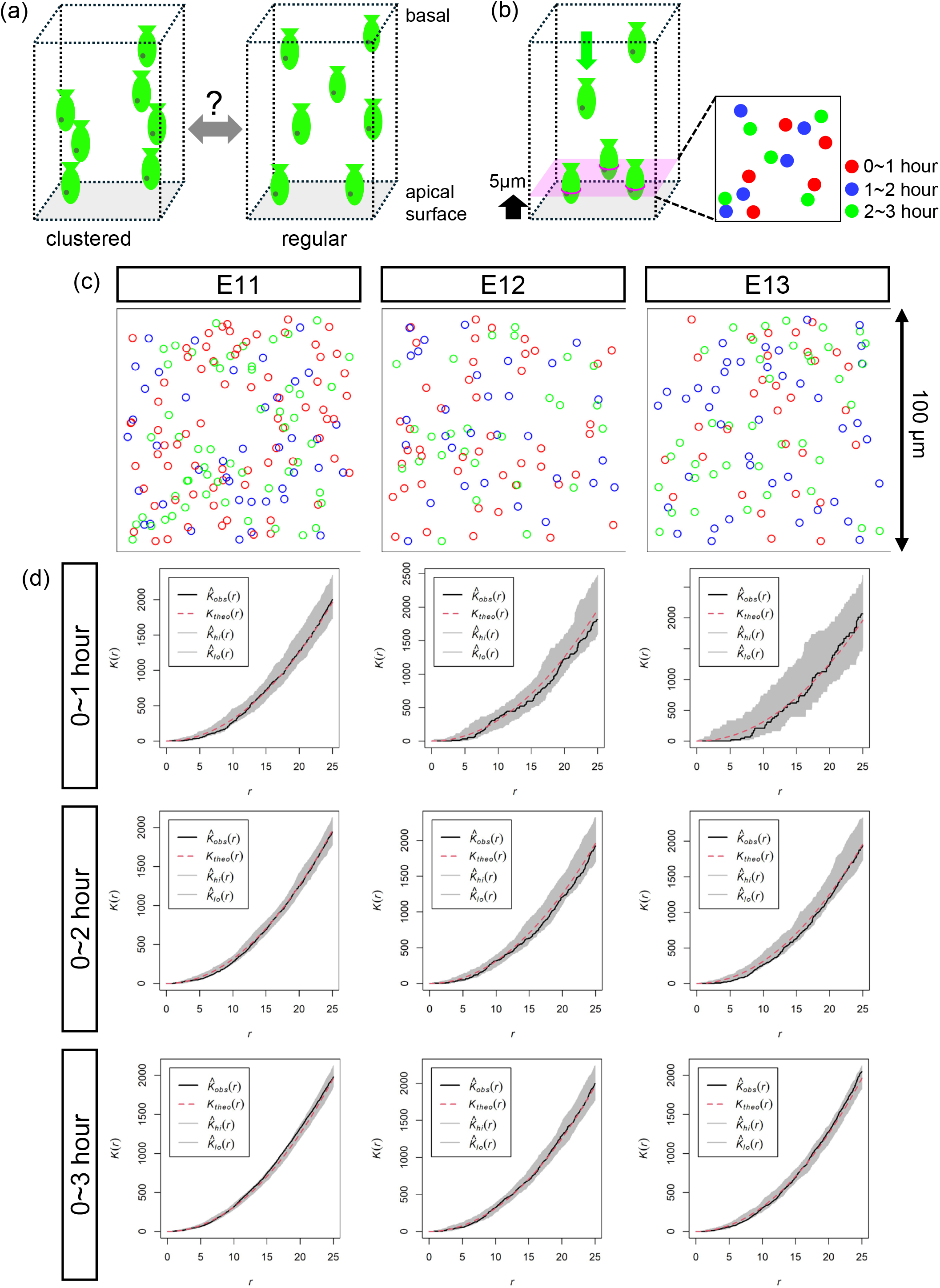
A comparison of the cumulative distribution patterns of mitoses was performed to infer the spatial distribution of apical nucleokinesis. (a) Schematic illustration of a pattern called “clustered”, in which the apically migrating nuclei and resulting mitoses are closely grouped, and another pattern called “regular”, in which the apically migrating nuclei and resulting mitoses are observed at a certain distance from each other. (b) Schematic illustration of the cumulative analysis of the horizontal distribution of mitoses. Three coordinates (x, y, t) were live recorded for each mitosis at 5 mm from the apical surface and were pseudocolored depending on the time of observation. (c) Examples of the recorded apical mitoses and premitotic nucleokinesis, superimposed on cumulative observations for 3 hours at E11, E12, or E13. (d) Graph comparing the distribution pattern plot, which is calculated via Ripley’s K function for each time duration, between E11, E12, and E13. Black line, calculated plot. The red broken line represents the theoretical random line (Poisson distribution). The gray zone indicates the 95% range of the simulated random pattern.

To this end, we analyzed the horizontal displacement of all nuclei at 30 μm (as the stage-dependent densified position) and 60 μm (as the stage-dependent non-densified position) from the apical surface by calculating the mean square displacement (MSD). MSD is useful for evaluating various types of movements at the molecular and cellular levels, including the apicobasal IKNM (Norden et al., 2009; Okamoto et al., 2013). Therefore, at a given horizontal plane, obtaining the MSD for all the nuclei at a given apicobasal level allows us to assess mainly the lateral mobility of the non-G2-phase nuclei. As shown in the graph depicting the MSD plotted versus the elapsed time at 30 μm from the apical surface (Figure 4c), the horizontal displacement of nuclei showed no directional preferences (thus judged to be random) at E11, E12, and E13. In the MSD–time interval graph, the initial slope is proportional to the diffusion coefficient, which represents how far each displacement occurred in unit time. At each depth, the diffusion coefficient significantly decreased in a stage-dependent manner: the diffusion coefficient at 30 μm depth was 0.175 ± 0.011 mm^2^/min at E11, 0.093 ± 0.004 mm^2^/min at E12, and 0.065 ± 0.003 mm^2^/min at E13 (mean ± se), whereas the diffusion coefficient at 60 μm depth was 0.129 ± 0.005 mm^2^/min at E11, 0.081 ± 0.003 mm^2^/min at E12, and 0.065 ± 0.003 mm^2^/min at E13 (mean ± se). The diffusion coefficient was smaller at 60 μm depth than at 30 μm depth at E11 and E12 (p < 0.001). At E13, however, the diffusion coefficient was comparable between 30 and 60 μm. The spatiotemporal changes found in the horizontally obtained diffusion coefficient within the NE were consistent with the changes in intra-NE nuclear density (Figure 2) and those in the live observed apical nucleokinesis (Figure 1). These results suggest that the degree to which a given horizontal plane of the NE flexibly drifts laterally, which can be permissive to apical nucleokinesis, decreases from E11 to E13.

### Apical nucleokinesis was assessed to occur in a horizontally random pattern

The aforementioned stage-dependent difference in the horizontal mobility of the NE prompted us to further ask whether the apical nucleokinesis flows are distributed in a certain pattern within the horizontal planes of the NE. One pattern is that the flows of apical nucleokinesis are mutually separated from each other (“regular”). Another pattern is that apical nucleokinesis flows are closely associated with each other (“aggregated” or “clustered”). We sought to determine whether the NE becoming less permissive to apical nucleokinesis (less flexibly drifting laterally) between E11 and E13 affects the horizontal distribution of the apical nucleokinesis flows. Using movies obtained through in toto recording at the level of 5 mm from the apical surface, we performed a cumulative analysis on the position of the mitoses. At a given 0.1 mm × 0.1 mm field, mitotic events during a period of 3 hours were superimposed. Through cumulative plotting of both the currently occurring mitoses and those that had occurred within the last 3 hours in the same xy field, the horizontal distribution of mitoses could be analyzed more thoroughly than by simply using a snapshot at one time point.

To statistically determine to which of the regular, clustered, or random patterns the cumulatively recorded cytokinesis positions at E11, E12, or E13 belong, we used Ripley’s K-function, which is widely used for multidistance spatial cluster analysis (Jafari Mamaghani et al., 2010; Barry-Carroll et al., 2023; Dayao et al., 2023). Ripley’s K function is defined as the expected number of additional points that lie within a radius (*r*) set for each point in the pattern (Ripley, 1987). It statistically compares a point distribution observed (e.g., distribution of proteins in the membrane microdomain or mitoses on the apical surface) with a random distribution mathematically simulated (Poisson configuration). When K(*r*) of an observed pattern is greater than that of the simulated random pattern, the observed pattern is judged as clustered, whereas another observed pattern showing a K(*r*) smaller than that of the simulated random pattern is judged to be regular or dispersed. Unexpectedly, the distribution pattern of mitoses obtained through the cumulative analysis of in toto recorded movies suggests that despite the stage-dependent reduction in the horizontal mobility of the NE from E11 to E13, as revealed by MSD analysis, the horizontal distribution of NPC mitoses as the result of apical nucleokinesis within the NE was judged to be consistently random throughout the period from E11 to E13.

## Discussion

### Cell intrinsic and extrinsic aspects of apical nucleokinesis regulation

In the present study, we comprehensively analyzed the apicalward IKNM behaviors of E11-E13 mouse telencephalic NEs via multiple imaging approaches, including cellular visualization. Comparison of the velocity of apical nucleokinesis exhibited by G2-phase NPCs and the density of nuclei in an intra-NE subzone where the apically migratory nuclei need to pass through allowed us to establish a model in which the apical nucleokinesis decelerates under physical influences from the densification of the surrounding intra-NE space. This model was supported by our Shh overexpression experiment in vivo, in which pulse-chase analyses revealed that fewer nuclei corresponding to those of G2-phase cells were observed in a subzone of the nuclear densified NE than in the same subzone of the control NE. Interestingly, a previous comparative study on the telencephalic NE between a mouse (E13) and a ferret (E29, corresponding to the mouse E13) (Okamoto et al., 2014) revealed histological differences between the ferret and mouse NE (i.e., the ferret NE is apicobasally thicker with a greater nuclear density at the 60-80 mm level than the mouse NE). Since these differences resemble those between the shh-OE NE and the control NE that we observed in this study, the fact that apical nucleokinesis was slower in ferrets than in mice (Okamoto et al., 2014) is also in accordance with the model of nuclear density-dependent deceleration of apical nucleokinesis. As another previous study in Drosophila also revealed that cellular densification affects IKNM speed (Kirkland et al., 2020), the susceptibility of IKNM to external physical factors may be conserved across species. However, the density of nuclei in the abovementioned ferret NE is lower at the 16-μm intra-NE level (Okamoto et al., 2014), suggesting that external factors, including nuclear density, may not solely explain the velocity of apical IKNM. At this point, we do not know how NPCs’ ability to sense/recognize external obstacles or physical restrictions leads to or interacts with intracellular molecular mechanisms. It is interesting to investigate whether the two different dynein pathways, i.e., the RanBP2-BicD2 pathway working at > 30 μm intra-NE levels and the Nup133-CENP-F pathway working more apically (Hu et al., 2013), differentially participate in the alterations in apical nucleokinesis that we observed in an NE depth-dependent manner.

### Lessons from the four-dimensional inferred randomness in the horizontal distribution of apical nucleokinesis

En face or horizontal sectional views at a given timepoint provide two-dimensional information about the x‒y position of the apically undergoing mitoses. Our cumulative analysis (for 3 hours) added a temporal dimension to this analysis, thereby also partly superimposing the trajectory of apical nucleokinesis, thus adding another (z) axis. Based on the stage-dependent change in the horizontal mobility of NE subzones caused by intra-NE nuclear densification, we expected that the horizontal distribution pattern of the apical mitoses (as a readout of the apical nuclear flows) may also change in a stage-dependent manner. Our statistical analysis using Ripley’s K-function, however, did not reveal a drastic change, i.e., conversions from any of the three candidates (clustered, regular, and random) to others. Nevertheless, the fact that the distribution of the apical mitoses was judged to be “random” throughout from E11 to E13 is interesting. This consistent preference of the random distribution may represent the robustness of the collective IKNM system of the telencephalic NE. It will be interesting to examine whether this is also the case for other brain regions and whether there are species-specific differences at this point. The “random” horizontal placement of IKNM trajectories is accomplished through the asynchronization of IKNM between neighboring cells. One likely mechanism that may underlie such asynchronized IKNM is seen in the initial phase of basal nucleokinesis; pair-generated daughter cells move their nuclei sequentially via a basal process asymmetrically inherited from their parent NPC (Okamoto et al., 2013). Intercellular mechanical interactions that regulate cell cycle progression have also been explored (Hacht et al., 2022). Therefore, the random pattern may represent the results of successful execution of multiple community-level mechanisms involving mechanosensing and/or mechanochemical coupling. Pathology, such as overacceleration of nucleokinesis or inability to sense, could affect the entire nucleokinesis pattern, perhaps generating nonrandom patterns.

## Author contributions

T.S. and T.M. designed the study. T.S. performed all the experiments and data analysis. T.S. and T.M. wrote the manuscript.

## Acknowledgments

We thank Makoto Masaoka and Namiko Noguchi (Department of Anatomy and Cell Biology, Nagoya University Graduate School of Medicine) for their technical assistance. The authors wish to acknowledge the Division for Medical Research Engineering, Nagoya University Graduate School of Medicine, for usage and technical support of the multiphoton excitation microscopy system [A1RMP (Nikon, Tokyo, Japan)]. T.S. was supported by the Nagoya University CIBoG WISE program from MEXT. This work was supported by JSPS 24K02205 (T.M.), MEXT 23H04306 (T.M.), JSPS 23K18339 (T.M.), MEXT 21H00363 (T.M.), and JSPS 21H02656 (T.M.).

